# Proteomic analysis of papillary thyroid carcinoma in the context of Hashimoto’s thyroiditis

**DOI:** 10.1101/2024.09.19.614028

**Authors:** Hui Zhou, Hui Sun, Muhammad Asad Iqbal, Ziou Zhao, Jie Hou, Xian Wang, Donggang Pan

**Affiliations:** Department of Ultrasound, Affiliated People’s Hospital of Jiangsu University, Zhenjiang, China; Department of Pathology, Affiliated People’s Hospital of Jiangsu University, Zhenjiang, China; School of Medicine, Jiangsu University, Zhenjiang, Jiangsu, 212013, China; Department of Radiology, Affiliated People’s Hospital of Jiangsu University, Zhenjiang, China

**Keywords:** papillary thyroid carcinoma, Hashimoto’s thyroiditis, proteomics, differential proteins, specific molecular markers

## Abstract

**Objective:** To analyze the proteomic characteristics of papillary thyroid carcinoma in the context of Hashimoto’s thyroiditis by mass spectrometry and screen the corresponding differential proteins.

**Methods:** Postoperative paraffin specimens of 3 patients with Hashimoto’s thyroiditis combined with papillary thyroid carcinoma and 3 patients with Hashimoto’s thyroiditis and benign nodules were collected. The differential proteins were obtained by qualitative and quantitative analysis using Q ExactiveTM mass spectrometer. Functional enrichment cluster analysis and protein interaction analysis were then performed on these differential proteins. Finally, key proteins were screened out and immunohistochemical verification was performed.

**Results:** 72 up-regulated differentially expressed proteins and 21 down-regulated differentially expressed proteins were screened out. THBS2 and COL12A1 were further identified by bioinformatics analysis. After immunohistochemical verification, it was concluded that COL12A1 can be used as a specific tumor marker.

**Conclusion:** There are differentially expressed proteins between the Hashimoto’s thyroiditis combined with papillary thyroid carcinoma group and the Hashimoto’s thyroiditis combined with benign nodules group. The up-regulated expression of COL12A1 can become an auxiliary diagnostic marker for papillary thyroid carcinoma with Hashimoto’s thyroiditis.

## Introduction

Thyroid cancer is a common endocrine tumor that originates from follicular cells and neuroendocrine cells(1). The most common pathological type is papillary thyroid carcinoma (PTC), which accounts for about 80% of all cases, and its incidence rate is increasing year by year worldwide(2, 3). Hashimoto’s thyroiditis (HT) is a common chronic autoimmune disease that uses its own thyroid tissue as an antigen(4). Studies have shown that there is a certain correlation between the occurrence of thyroid cancer and HT(5). HT increases the risk of PTC, but has an inhibitory effect on the metastasis, progression, and recurrence of thyroid cancer(5, 6). Currently, ultrasound examination is the preferred imaging examination for diagnosing thyroid cancer(7). However, since HT causes the thyroid gland to show uneven echoes on ultrasound images, the diagnostic accuracy of ultrasound for thyroid cancer with HT is significantly reduced. A study reported(8) that the detection rate of preoperative ultrasound for tumors in the context of diffuse thyroid lesions was 68.21%, and the misdiagnosis rate was 31.79%. Timely and accurate diagnosis of whether HT is accompanied by a malignant tumor or a benign nodule is of great significance to the patient’s treatment selection and prognosis.

The composition and interaction of proteins constitute multiple processes of human physiological activities, participate in the structure of human tissues and organs, and form the various functions and regulation of tissues and organs(9). Therefore, the occurrence and development of diseases are often accompanied by changes in the quantity and quality of proteins. Under different physiological or pathological conditions, by classifying and identifying proteins, analyzing the difference in protein expression levels, and studying the interaction patterns and characteristics between proteins, the mechanism of the disease can be revealed. With the development of molecular biology technology, the diagnosis of thyroid cancer is moving towards molecular diagnosis(10).

Mass spectrometry is an important tool for identifying, characterizing and quantifying proteins. It has the characteristics of high throughput and large scale(11), which has promoted the rapid development of proteomics and made it possible to use proteomics in the diagnosis of thyroid cancer. Therefore, this study conducted TMT-labeled qualitative and quantitative proteomics analysis on paraffin-embedded tissue specimens of HT combined with PTC and HT with benign nodules after surgery to explore the specific differential proteins between the two, and then screened out protein markers that can be used for auxiliary diagnosis of HT combined with PTC, to find its possible mechanism of occurrence and provide a theoretical basis for the development of targeted diagnosis and treatment.

## Materials and methods

### Research subjects

This study was conducted in accordance with the ethical standards outlined in the Declaration of Helsinki. Ethical approval for this research was obtained from the Ethics Committee of Jiangsu University Affiliated People’s Hospital, with the approval number K-20230119-W. All participants provided written informed consent prior to their inclusion in the study, and their confidentiality was strictly maintained throughout the research process.

This study collected 3 patients with HT combined with PTC and 3 patients with HT and benign nodules admitted to the Affiliated People’s Hospital of Jiangsu University from 1-9-2020 to 31-11-2021. The inclusion criteria were: (1) surgery in our hospital and a clear diagnosis by pathology; (2) the surgically removed tissue could be made into formalin-fixed paraffin samples. Exclusion criteria: patients with underlying diseases such as hypertension, diabetes or other liver and kidney dysfunction.

Paraffin section specimens of 18 patients with HT combined with PTC and 17 patients with HT and benign nodules pathologically confirmed in the Affiliated People’s Hospital of Jiangsu University from 1-9-2020 to 31-1-2023 were collected. This study has been approved by the Ethics Committee of the Affiliated People’s Hospital of Jiangsu University, and all patients have signed informed consent.

### Research methods

Three samples of HT combined with PTC and three samples of HT with benign nodules were cryogenically ground into powder and quickly transferred to a centrifuge tube pre-cooled with liquid nitrogen. An appropriate amount of PASP protein lysis buffer (100mM ammonium bicarbonate, 8M urea, pH=8) was added, and the mixture was shaken and mixed. The samples were fully lysed by ultra-sonication in an ice-water bath for 5 minutes. The samples were centrifuged at 4°C and 12000g for 15 minutes. The supernatant was added with 10mM DTT and reacted at 56°C for 1 hour. Then, sufficient IAM was added and reacted at room temperature in the dark for 1 hour. Four volumes of -20°C pre-cooled acetone were added and precipitated at -20°C for at least 2 hours. The samples were centrifuged at 4°C and 12000g for 15 minutes, and the precipitate was collected. Then add 1 mL -20°C pre-cooled acetone to resuspended and wash the precipitate, centrifuge at 4°C and 12000g for 15 min, collect the precipitate, air-dry, and add an appropriate amount of protein dissolving solution (8 M urea, 100 mM TEAB, pH=8.5) to dissolve the protein precipitate.

Take the protein sample, add DB protein dissolving solution (8M urea, 100 mM TEAB, pH=8.5) to make up the volume to 100μL, add trypsin and 100 mM TEAB buffer, mix well and digest at 37°C for 4h, then add trypsin and CaCl2 for digestion overnight. Add formic acid to adjust the pH to less than 3, mix well and centrifuge at room temperature and 12000g for 5 min, take the supernatant and slowly pass it through the C18 desalting column, then use the cleaning solution (0.1% formic acid, 3% acetonitrile) to wash three times continuously, then add an appropriate amount of eluent (0.1% formic acid, 70% acetonitrile), collect the filtrate, and freeze-dry. Add 100 μL 0.1 M TEAB buffer to reconstitute, and add 41 μL TMT labeling reagent dissolved in acetonitrile, invert and mix at room temperature for 2 hours. Then add ammonia water at a final concentration of 8% to terminate the reaction, take an equal volume of labeled samples, mix, desalt and freeze-dry.

Prepare mobile phase A (2% acetonitrile, 98% water, ammonia water adjusted to pH=10) and B (98% acetonitrile, 2% water). Dissolve the mixed freeze-dried powder with A solution and centrifuge at 12000 g for 10 min at room temperature. Use L-3000 HPLC system, the chromatographic column is Waters BEH C18 (4.6×250 mm, 5 μm), the column temperature is set to 45°C, collect 1 tube per minute, merge into 10 fractions, and add 0.1% formic acid to each fraction after freeze-drying.

Prepare mobile phase A (100% water, 0.1% formic acid) and B (80% acetonitrile, 0.1% formic acid). 1 μg of the supernatant of each fraction was injected for LC/MS detection. The EASY-nLCTM 1200 nanoliter UHPLC system was used, with a homemade precolumn (4.5 cm × 75 μm, 3 μm) and a homemade analytical column (15 cm × 150 μm, 1.9 μm). A Q ExactiveTM series mass spectrometer and Nanospray FlexTM (ESI) ion source were used. The ion spray voltage was set to 2.3 kV, the ion transfer tube temperature was set to 320°C, the mass spectrometer adopted data-dependent acquisition mode, the full scan range of the mass spectrometer was m/z 350-1500, the primary mass spectrometer resolution was set to 60000 (200 m/z), the C-trap maximum capacity was 3×106, and the C-trap maximum injection time was 20 ms; the parent ions with the TOP 40 ion intensities in the full scan were selected and fragmented using the high energy collision fragmentation (HCD) method for secondary mass spectrometry detection. The secondary mass spectrometry resolution was set to 45000 (200 m/z), the C-trap maximum capacity was 5×104, the C-trap maximum injection time was 86 ms, the peptide fragmentation collision energy was set to 32%, the threshold intensity was set to 1.2×105, and the dynamic exclusion range was set to 20 s to generate the mass spectrometry detection raw data (. raw). The original data was searched for the result spectrum of each run using the search software Proteome Discoverer2.4. The search parameters were set as follows: the mass tolerance of the precursor ion was 10ppm, and the mass tolerance of the fragment ion was 0.02Da. The fixed modification was alkylation of cysteine, the variable modification was methionine oxidation and TMT tag modification, the N-terminus was acetylation and TMT tag modification, and a maximum of 2 missed cleavage sites were allowed.

In order to improve the quality of the analysis results, the PD2.2 software further filtered the search results: the peptides (Peptide Spectrum Matches, PSMs) with a credibility of more than 99% were credible PSMs, and the proteins containing at least one unique peptide were credible proteins. Only credible peptides and proteins were retained, and FDR verification was performed to remove peptides and proteins with FDR greater than 1%.

### Bioinformatics analysis

The data after mass spectrometry analysis were analyzed for differences, with P < 0.05 and fold charge (FC) greater than 1.2 as significantly upregulated differential proteins, and less than 1/1.2 as significantly downregulated differential proteins, to identify the proteins with significant differences. The online database metascape (https://metascape.org/) was used to perform pathway enrichment analysis on the identified differential proteins, and cytoscape was used to construct a protein interaction network. Based on MCODE clustering and cytohubba algorithms, key proteins were screened, and the online database GeneMANIA (http://genemania.org/) was used to predict the gene functions of the key proteins obtained. Finally, GSEA enrichment analysis was used to find the most significantly enriched pathways and confirm the key proteins.

### Immunohistochemistry

After dewaxing the tissue sections of 35 cases, antigen repair was performed, and an appropriate amount of primary antibody working solution diluted with PBS was added to the tissue for antigen-primary antibody reaction, and then an appropriate amount of ready-to-use rapid immunohistochemistry MaxVisionTM HRP secondary antibody corresponding to the primary antibody was added for primary antibody-secondary antibody reaction. After color development with 3,3-diaminobenzidine (DAB), the cell nucleus was counterstained, the slides were dehydrated and transparently sealed, and the results were scanned with a scanner for judgment.

### Statistical analysis

SPSS 25.0 statistical software was used for data analysis, and pairwise comparisons were performed using t-test (measurement data) and χ2 test (count data) or Mann-Whitney U test. Fisher’s exact probability method was used for subgroup analysis. All tests were two-sided tests, and P < 0.05 was the significance judgment standard.

## Results

### Basic information of research subjects

This study included 3 patients with HT combined with PTC (tumor group) and 3 patients with HT and benign nodules (control group), as well as 18 patients with HT combined with PTC (PTC group) and 17 patients with HT and benign nodules (benign nodule group) for immunohistochemical verification, see Table 1. The clinical data of the PTC group and the benign nodule group were statistically compared and analyzed, and there was no statistically significant difference in laboratory tests between the two groups of patients, see Table 2.

**Table 1.**
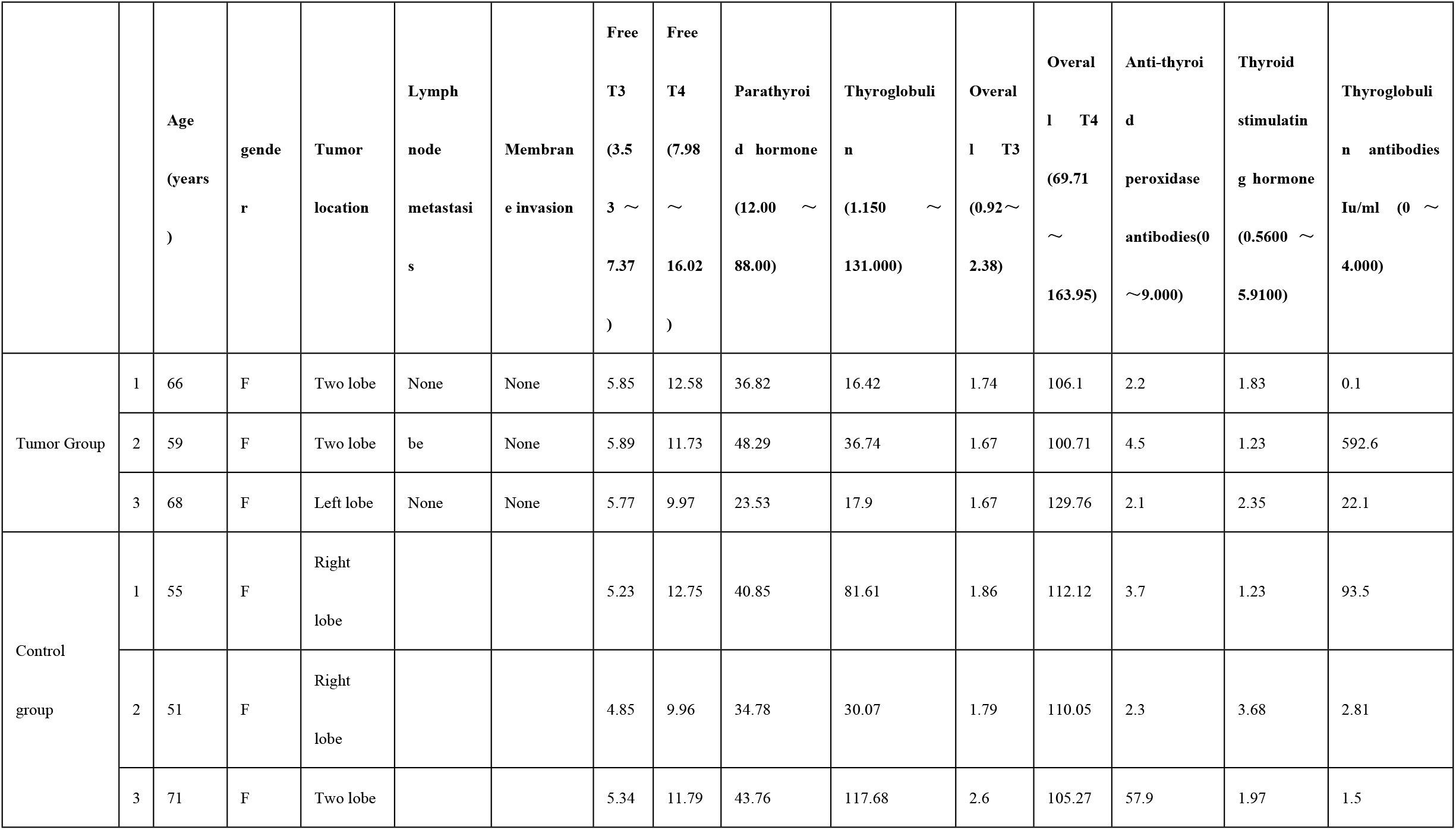
Comparison of clinical data between tumor group and control group.

**Table 2.**
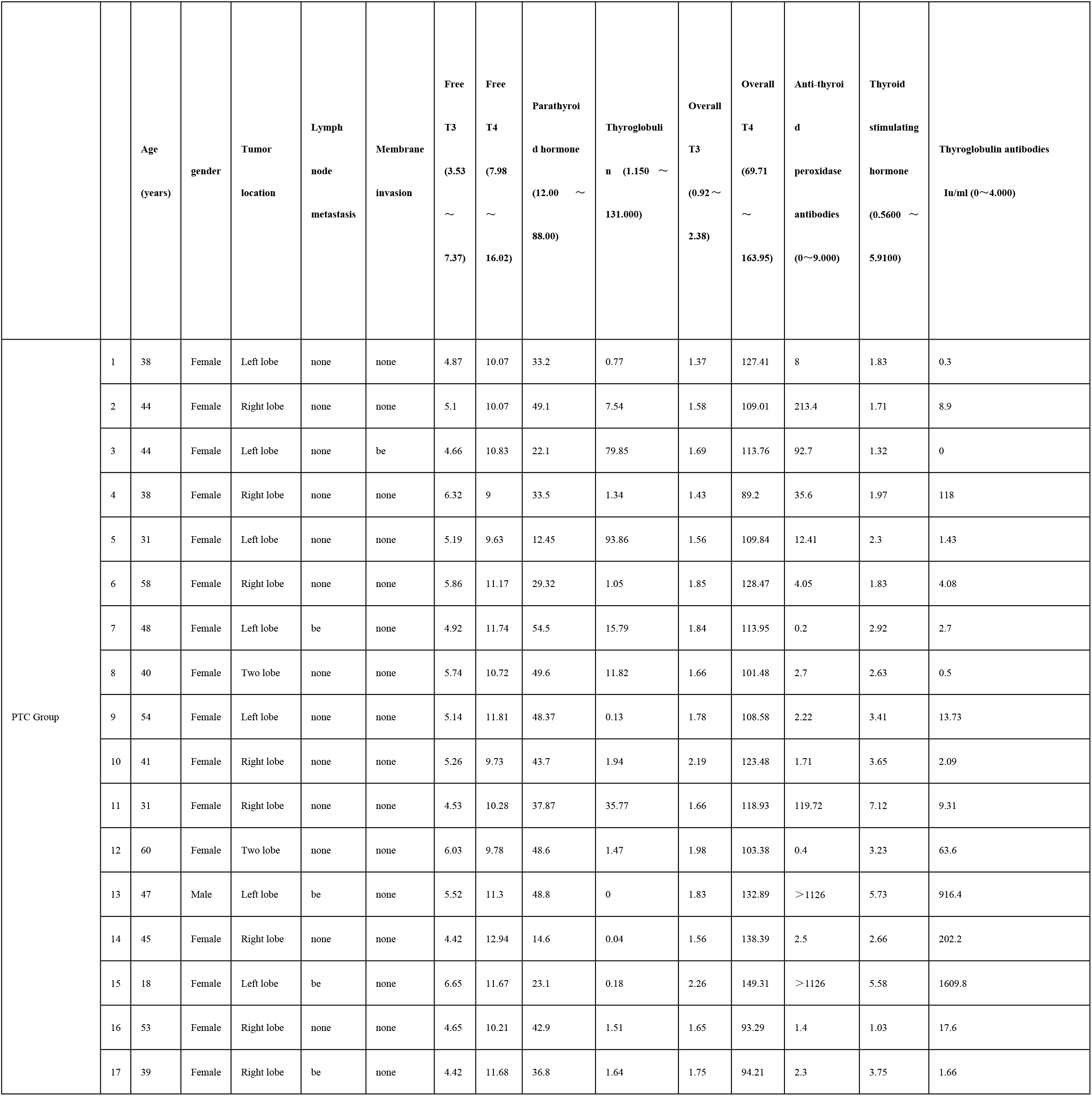
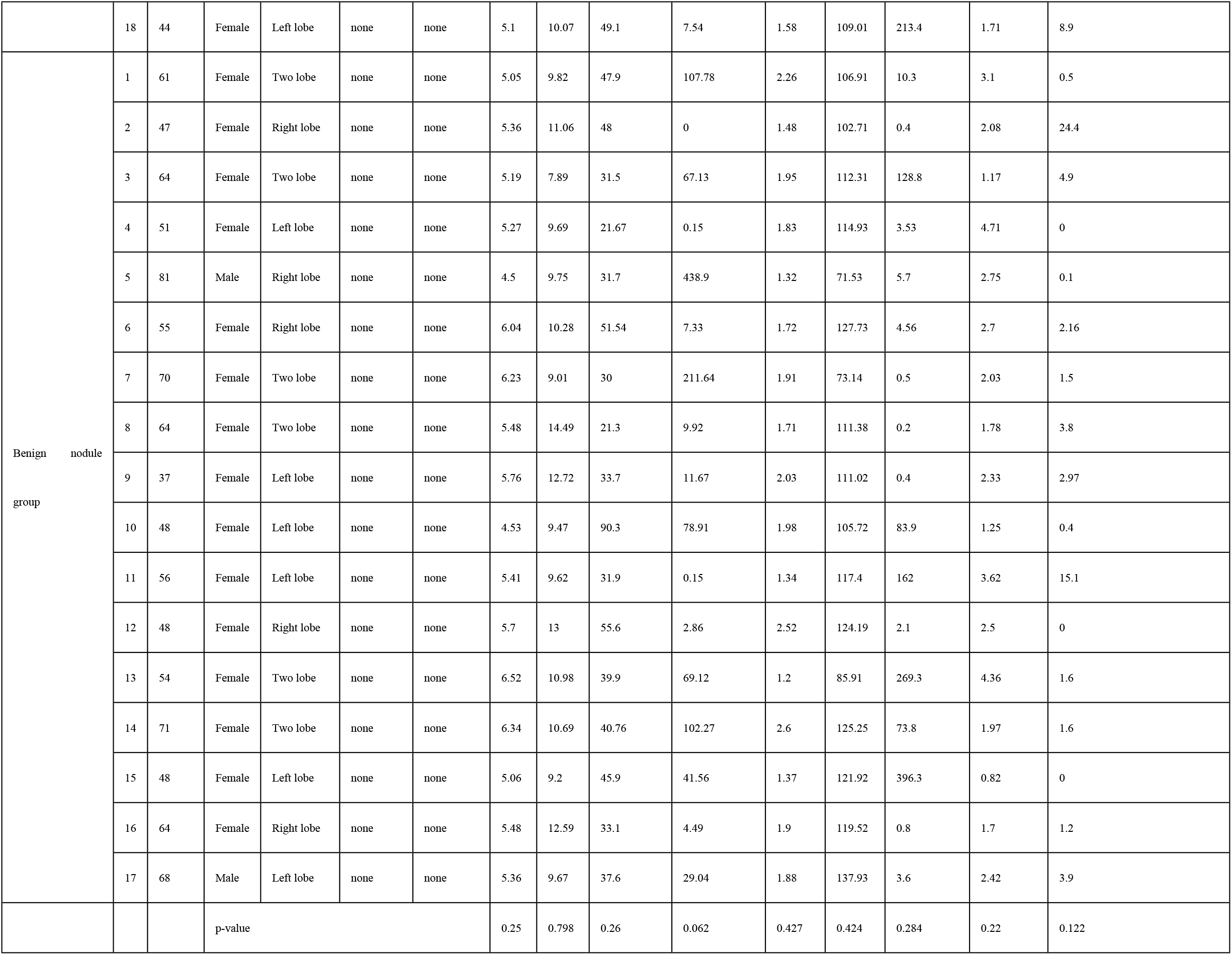
Clinical data comparison between PTC group and benign nodule group.

### Protein differential analysis results

Through mass spectrometry analysis, we obtained 59,367 valid spectra and identified 3,981 proteins, of which 3,934 were quantifiable. PCA analysis based on quantifiable proteins obtained two components (Figure 1A). The hierarchical clustering heat map also showed that there were significant differences in protein expression between PTC patients with HT (HT-PTC) and benign thyroid nodules with HT (HT-BTN) (Figure 1B). When the expression difference fold FC> 1.2 and Pvalue < 0.05 were met, 72 up-regulated differentially expressed proteins were screened, and when FC < 1/1.2 and Pvalue < 0.05 were met, 21 down-regulated differentially expressed proteins were screened(Figure1C).

**Figure 1.**
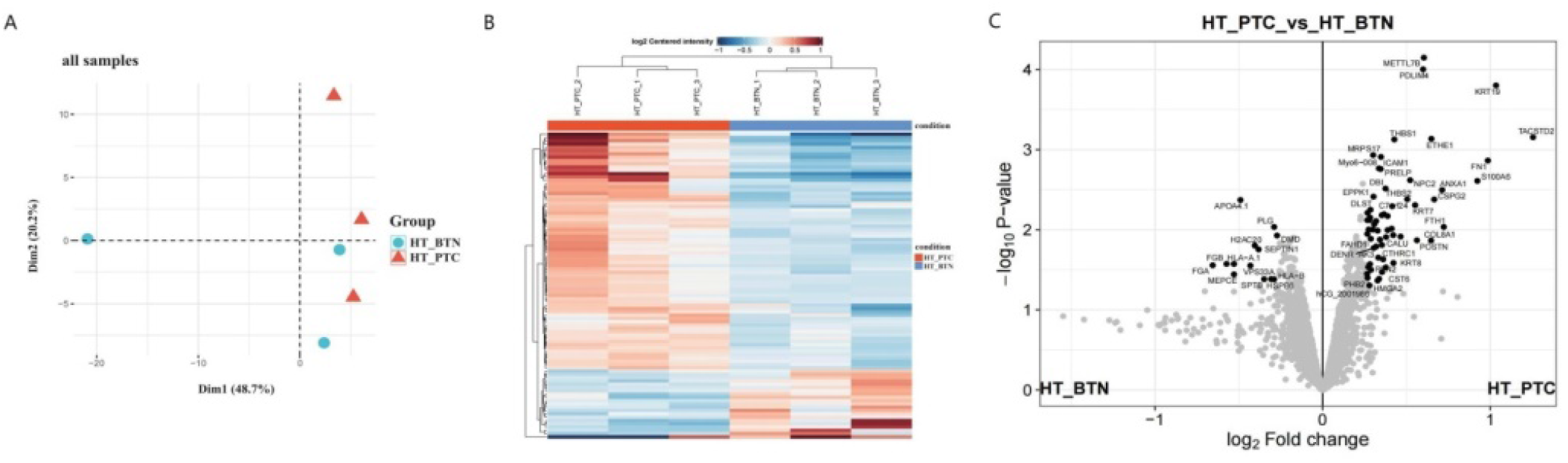
Analysis of differential protein expression between HT-PTC and HT-BTN groups. A: PCA analysis based on quantifiable proteins obtained two groups of components, with variance explanation rates of 48.7% and 20.2%, respectively; B: Hierarchical clustering results showed that there were significant differences in protein expression between HT-PTC and HT-BTN groups; C: A total of 93 differentially expressed proteins were identified in the HT-PTC and HT-BTN groups, of which 72 proteins were upregulated and 21 proteins were downregulated. HT-PTC: Patients with papillary thyroid carcinoma and Hashimoto’s thyroiditis; HT-BTN: Patients with benign thyroid nodules and Hashimoto’s thyroiditis.

### Bioinformatics analysis of differentially expressed proteins

Pathway enrichment analysis was performed on the differentially expressed proteins, and it was found that the up-regulated proteins mainly focused on the functions of Naba core matrix and metabolic diseases, supramolecular fibrous organization, regulation of epithelial cell migration, and extracellular matrix organization; the down-regulated proteins mainly focused on the functions of thrombosis, anticoagulation, and regulation of protein complex assembly (Figure 2).

**Figure 2.**
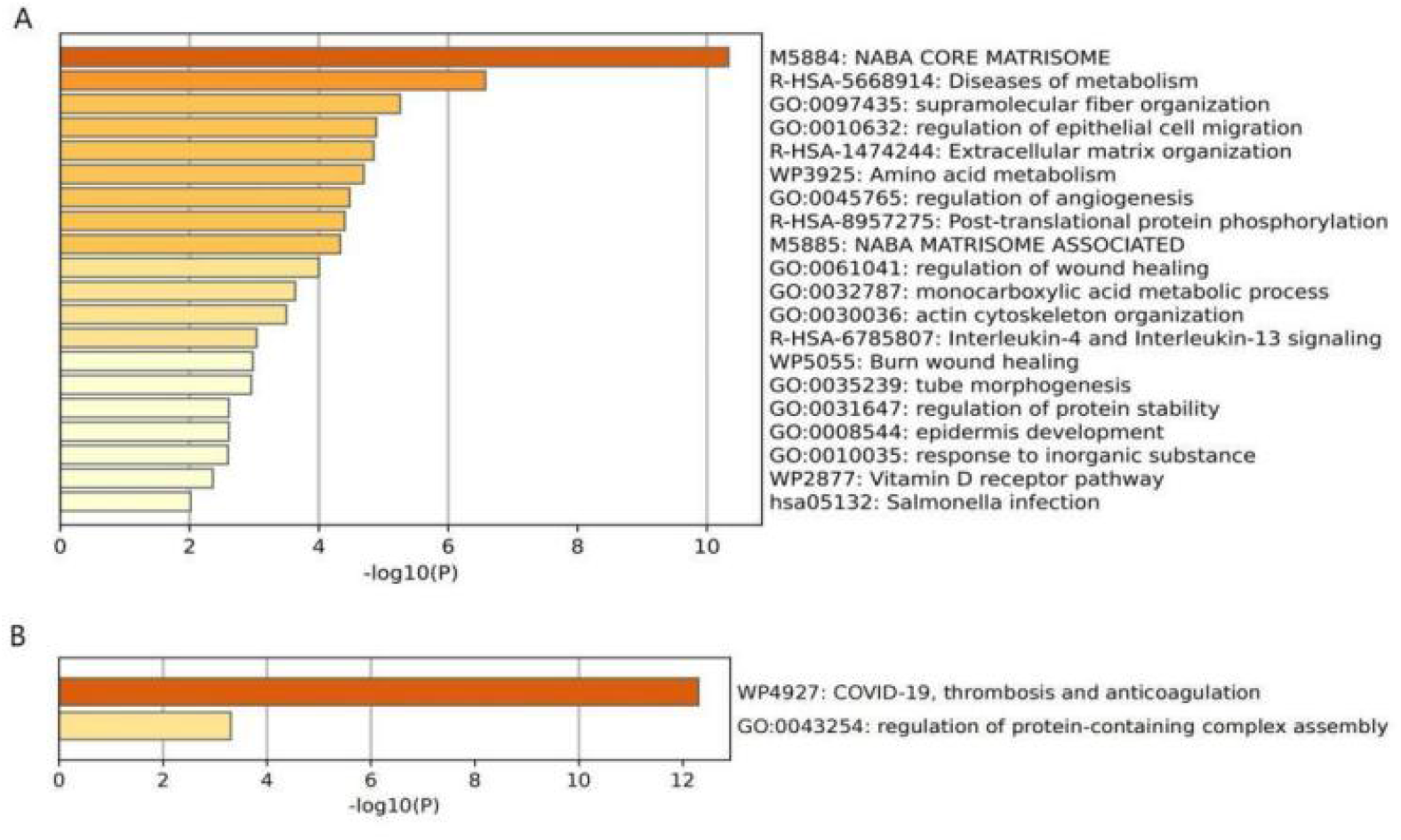
**Analysis of pathway enrichment of differentially expressed proteins A: Upregulated protein enrichment pathway; B: Downregulated protein enrichment pathway**

The protein-protein interaction network of 93 differentially expressed proteins was constructed using the STRING database, and a total of 39 interacting proteins were found, of which 32 were up-regulated differentially expressed proteins, such as THBS1, ANXA1, COL8A1, PLOD1, etc.; 7 were down-regulated differentially expressed proteins, such as PLG, FGB, etc. (Figure 3A). On this basis, MCODE (molecular complex detection) clustering and cytohubba were used to screen key proteins. By mining protein complexes and corresponding functional modules through MCODE, the protein network was analyzed and 4 modules were identified (Figure 3B). The CytoHubba plug-in in the Cytoscape software was used to discover the key targets and subnetworks of the complex network. The top 10 key proteins obtained by 5 different algorithms MCC, DMNC, MNC, Degree and EPC, respectively, can be obtained. Six overlapping proteins can be obtained, namely THBS2, COL12A1, PLG, COL8A1, VCAN, and ASPN (Figure 3C). Combining the MCODE clustering results and taking the intersection, five overlapping key proteins were obtained: THBS2, COL12A1, PLG, COL8A1, and ASPN.

**Figure 3.**
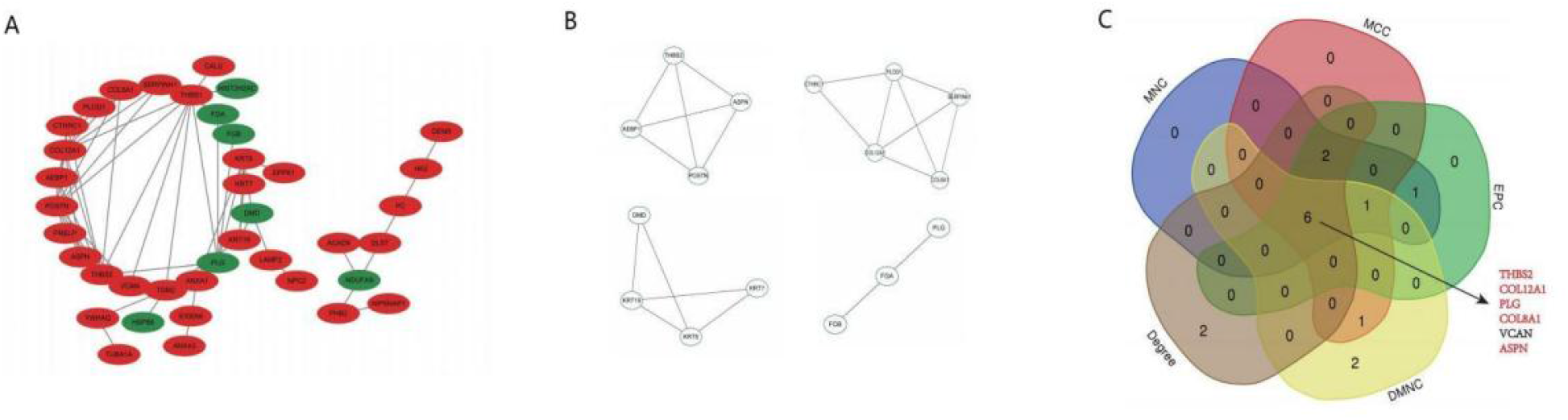
Screening of specific protein markers. A: In the PPI network, 32 proteins were upregulated (red) and 7 proteins were downregulated (green); B: MCODE clustering resulted in key proteins in 4 modules; C: 5 different algorithms of cytohubba and MCODE algorithm were used to screen key differential proteins.

Using the online database GeneMANIA to predict the gene functions of the five key proteins, 20 related proteins were obtained (Figure 4A). Functional enrichment analysis of the genes of the five key proteins and 20 related proteins was performed again, and it was found that the functions of these key proteins mainly focused on the regulation of coagulation function, extracellular matrix composition, PDGF signal transduction, and regulation of plasminogen activation (Figure 4B).

**Figure 4.**
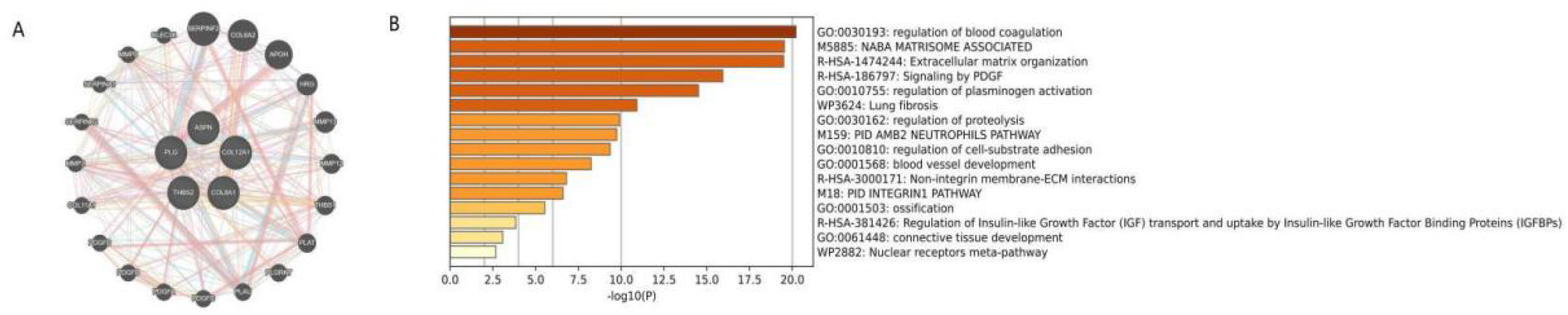
Function prediction of 5 key protein genes. **A: Protein interaction network with 5 key proteins as the core; B: Functional enrichment analysis of key proteins** **GSEA was used to conduct gene enrichment analysis of all proteins in the PTC group with HT**.

The pathway with the highest enrichment score was epithelial-mesenchymal transition (EMT) (NES= 1.90, q < 0.012), which contains 71 genes., some genes were significantly different between the two groups (Figure 5). Two of the key proteins overlapped with the aforementioned five, namely COL12A1 and THBS2, both of which were highly expressed in the tumor group (Figure 6).

**Figure 5.**
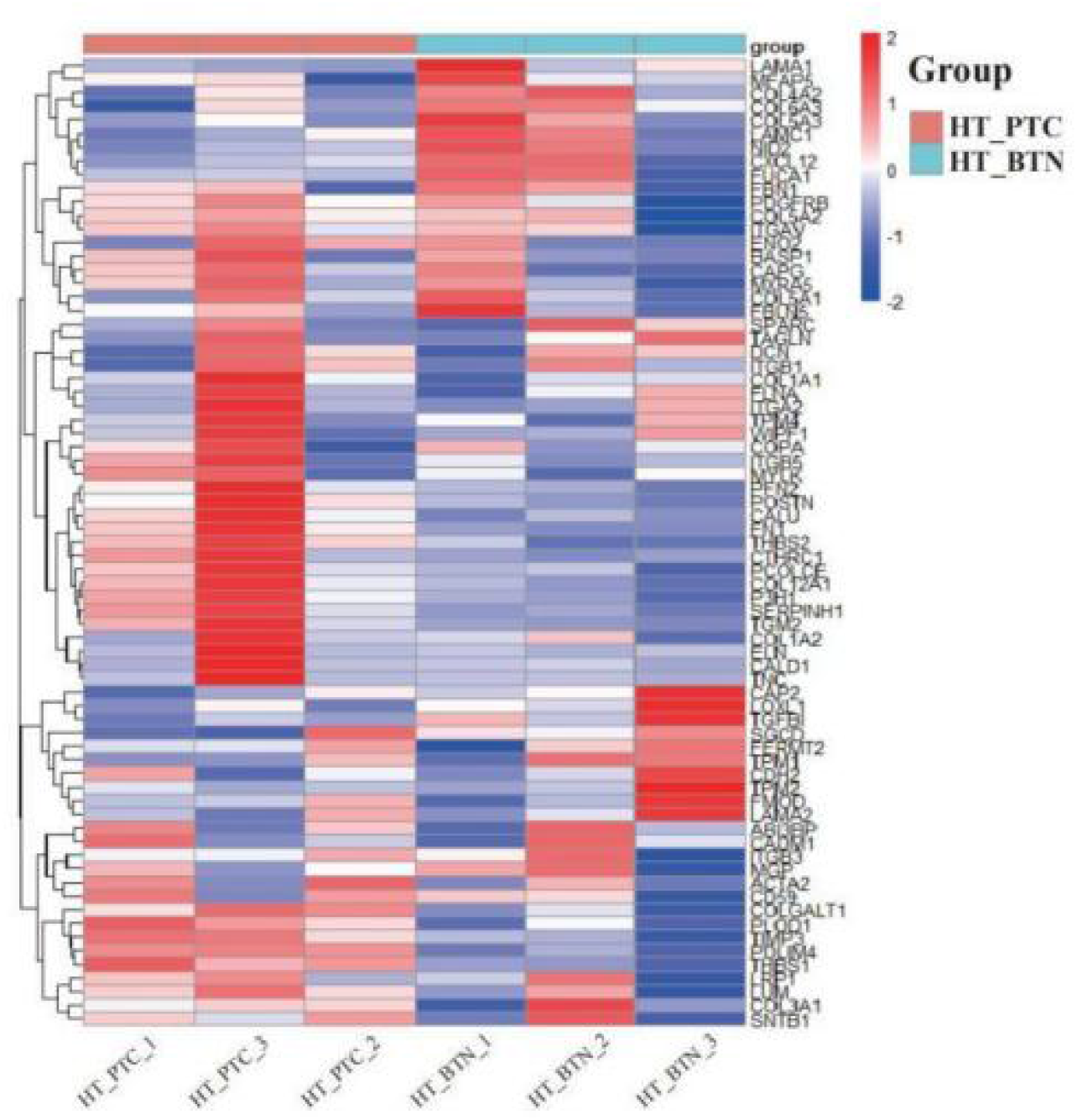
Analysis of differential expression of 71 genes related to the epithelial-mesenchymal transition (EMT) pathway between the HT-PTC and HT-BTN groups. **HT-PTC: patients with papillary thyroid carcinoma and Hashimoto’s thyroiditis; HT-BTN: patients with benign thyroid nodules and Hashimoto’s thyroiditis**

**Figure 6:**
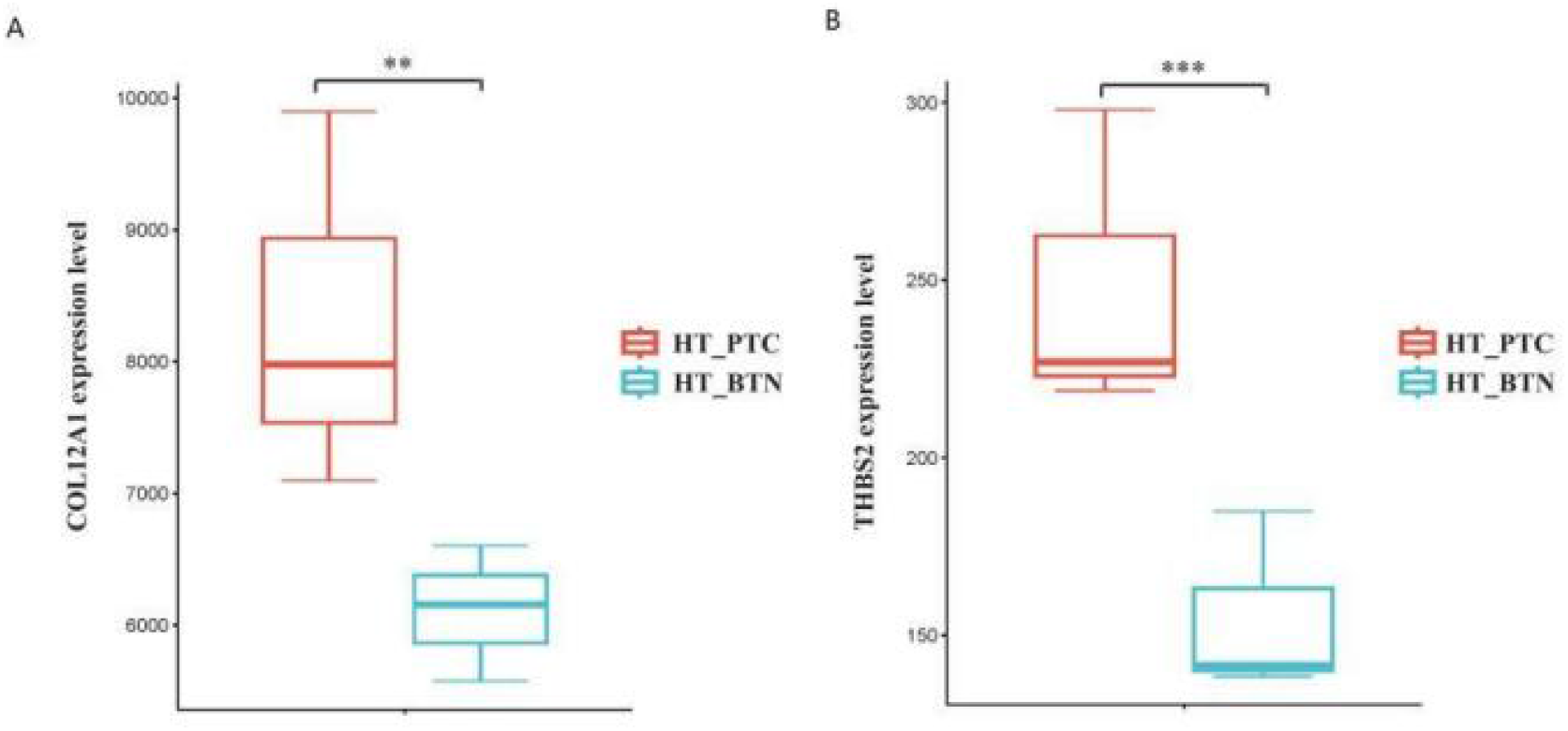
Box plot of COL12A1 and THBS2 expression between HT-PTC group and HT-BTN group. *** indicates p<0.001, ** indicates p<0.01. HT-PTC: Patients with papillary thyroid carcinoma and Hashimoto’s thyroiditis; HT-BTN: Patients with benign thyroid nodules and Hashimoto’s thyroiditis

**Figure 7:**
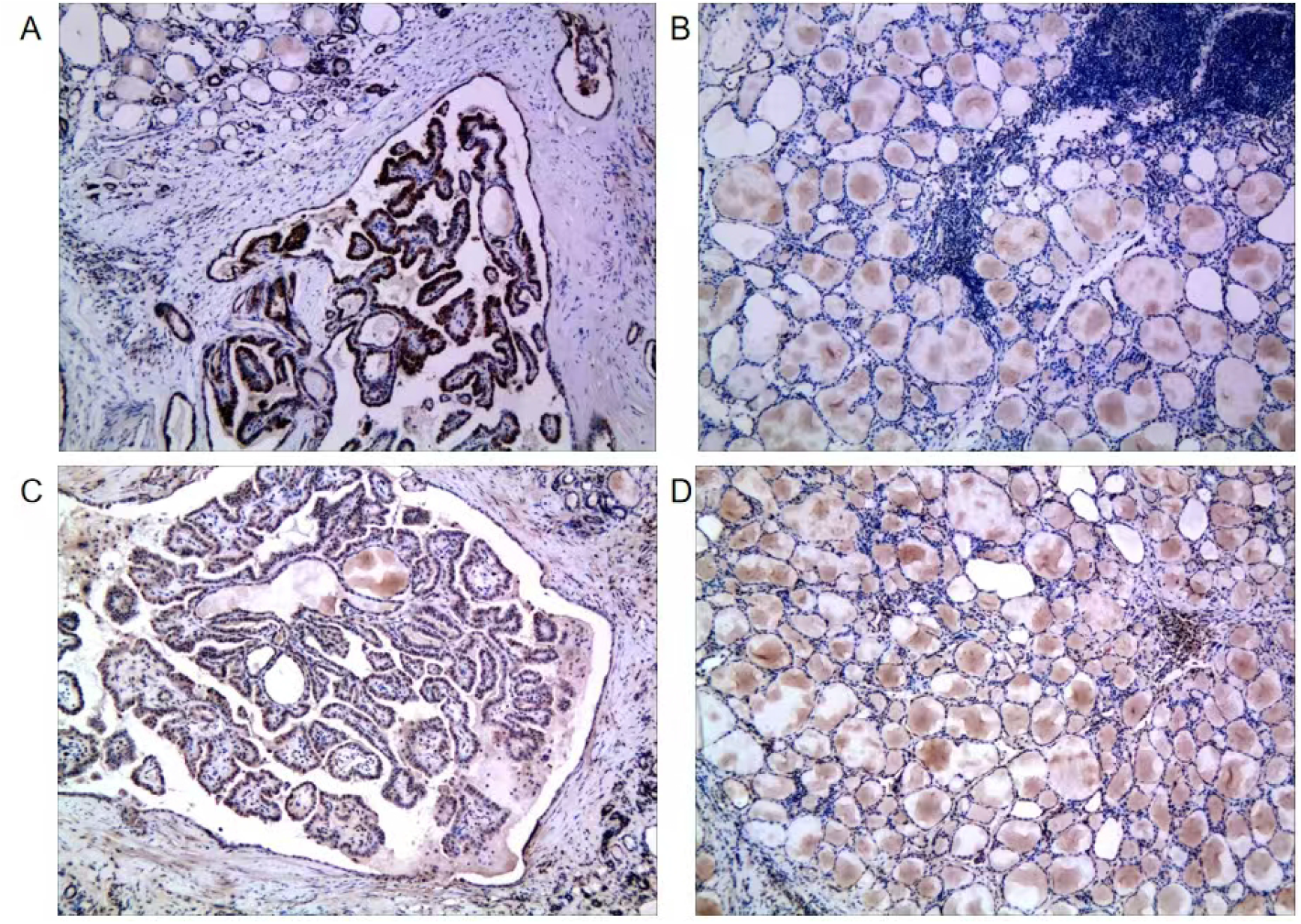
Immunohistochemical expression of COL12A1 and THBS2. Figure A: COL12A1 is positively expressed in HT with PTC tissue; Figure B: COL12A1 is negatively expressed in the thyroid tissue around the tumor; Figure C: THBS2 is positively expressed in HT with PTC tissue; Figure D: THBS2 is positively expressed in the thyroid tissue around the tumor

### Immunohistochemical expression

Among the 35 cases that underwent immunohistochemical staining, the positive rate of COL12A1 in the PTC group was 72.2% (13/18), and the positive rate in the benign nodule group was 29.4% (5/17); THBS2 was stained and positive in both the PTC group and the benign nodule group. The positive rate of COL12A1 in the PTC group was higher than that in the benign nodule group, and the difference was statistically significant (P<0.05).

## Discussion

In this study, compared with the HT with benign nodules group, 93 proteins with significantly differential expression levels were identified in the HT combined with PTC group, including 72 up-regulated differentially expressed proteins and 21 down-regulated differentially expressed proteins. After a series of bioinformatics analyses, including protein interaction analysis, MCODE clustering, and cytohubba screening of key proteins, two HT combined PTC-specific protein markers were finally obtained, COL12A1 and THBS2, both of which were found in the HT combined PTC group. Shows high expression. After immunohistochemistry verification, it was found that the positive rate of COL12A1 in the HT combined with PTC group was higher than that in the HT combined with benign nodules group, while THBS2 was positively expressed in both groups. Both COL12A1 and THBS2 are enriched in the epithelial-mesenchymal transition pathway. Epithelial-to-mesenchymal transition (EMT) is the process in which epithelial cells transform into mesenchymal cells and is involved in the malignant progression of tumors(12). Cell-cell and cell-extracellular matrix interactions are reshaped, resulting in reduced adhesion between epithelial cells and to the basement membrane, and the activation of new transcriptional programs to promote their transition to mesenchyme(13). Studies have shown that early-stage tumor cells are in an epithelioid state and gradually acquire more mesenchymal characteristics as the tumor progresses. Activation of EMT in tumor cells also induces tumors to be in a stem cell state, suggesting that EMT may be an integral part of the progression of all types of malignant tumors(14-17).

Type XII collagen encoded by COL12A1 is an important cell scaffold and supporting protein in the extracellular matrix. COL12A1 is found to be highly expressed in a variety of malignant tumors, such as pancreatic cancer(18), gastric cancer(19), and colorectal cancer(20). The occurrence and development of tumors are closely related to the proliferation of connective tissue and the remodeling of extracellular matrix(21, 22). Changes in cell polarity and loosening of intercellular adhesion in the tumor microenvironment are caused by changes in the composition and accumulation of extracellular matrix. occur, thereby promoting tumor growth, invasion and metastasis(23). Papanicolaou M et al(24) found that COL12A1 plays an important role in the metastasis of breast tumors. The increase in type XII collagen levels in primary tumors may strengthen matrix remodeling, thereby creating an environment that promotes metastatic spread and improves The aggressiveness of tumors is significantly correlated with tumor progression and poor prognosis. Jiang et al. (25) observed a significant upregulation of COL12A1 expression in gastric cancer. Clinicopathological analysis showed that elevated COL12A1 expression was positively correlated with tumor invasion, metastasis and clinical stage. In our study, COL12A1 was significantly up-regulated in patients with HT combined with PTC, while its expression was low in HT with benign nodules, indicating that COL12A1 may become an auxiliary diagnostic marker for HT combined with PTC.

The Tsp2 protein encoded by THBS2 is a member of the stromal cell Ca2+-binding glycoprotein family and is released by cells such as stromal fibroblasts, endothelial cells, and immune cells(26). It is a multifunctional glycoprotein with complex structural domains that can Promote the proliferation of microvascular endothelial cells and stimulate angiogenesis(27-29). It interacts with cell receptors and extracellular matrix proteins to promote cell adhesion, proliferation and apoptosis(30, 31), and its expression is up-regulated in various tumors such as pancreatic cancer, lung cancer, and gastric cancer(27, 32, 33). THBS2 has the potential to enhance EMT, and Zhang et al. found that THBS2 is related to tumor-related immune cell infiltration(34). Immunohistochemistry verification results in this study showed that THBS2 was positively expressed in both the HT combined with PTC group and the HT combined with benign nodules group, which may be related to the fact that HT causes the thyroid to undergo an autoimmune reaction, leading to inflammatory cell infiltration(35). On the other hand, since immunohistochemistry is a qualitative analysis of protein and cannot completely show the difference in protein expression, it cannot quantitatively distinguish the differential expression levels of THBS2 in HT combined with PTC and HT combined with benign nodules.

## Conclusion

This study used high-throughput protein spectrometry to analyze the proteomic characteristics of PTC tumors in the context of HT, and performed immunohistochemistry verification. It was found that COL12A1 was highly expressed in the HT combined with PTC patient group, and further explored the relationship between COL12A1 and PTC. It was finally determined that COL12A1 could be used as a target marker.

**Hui Zhou, Hui Sun and Muhammad Asad Iqbal contributed equally to this work**.

## Credit author statement

Hui Zhou, Hui Sun and Asad contributed equally to this study. Xian Wang, Hui Sun, Jie Hou,Donggang Pan contributed to the conception and design of the study. Jie Hou, Hui Zhou, organized the database. Asad, Ziou Zhao performed the statistical analysis. Hui Zhou wrote the first draft of the manuscript. Hui Sun and Xian Wang wrote sections of the manuscript. All authors contributed to the article and approved the submitted version.

